# Region-specific proteomic analysis of aging rhesus macaques following chronic glutamate-carboxypeptidase-II (GCPII) inhibition elucidates potential treatment strategies for sporadic Alzheimer’s disease

**DOI:** 10.1101/2025.11.07.687004

**Authors:** Alexandra D. Steigmeyer, Alexandria S. Battison, Iris R. Liebman, Christopher H. van Dyck, Barbara S. Slusher, Amy F. T. Arnsten, Stacy A. Malaker, Dibyadeep Datta

**Affiliations:** Department of Chemistry, Yale University, New Haven, CT, United States; Department of Psychiatry, Yale University School of Medicine, New Haven, CT, United States; Department of Neuroscience, Yale University School of Medicine, New Haven, CT, United States; Department of Neurology, Johns Hopkins University Drug Discovery, Johns Hopkins School of Medicine, Baltimore, MD, United States

**Keywords:** GCPII, calcium, macaque, mGluR3, proteomics, Alzheimer’s disease

## Abstract

Sporadic Alzheimer’s disease (sAD) lacks effective preventive therapies, underscoring the need to target pathogenic drivers. Aberrant calcium signaling is an established early event in sAD pathogenesis that is closely linked to neuroinflammation. Aged rhesus macaques are predominantly APOE-ε4 homozygotes and naturally exhibit cognitive decline, calcium dysregulation, amyloid deposition, and tau pathology, which allows for a translationally relevant animal model. We previously identified an evolutionarily expanded role for postsynaptic type 3 metabotropic glutamate receptors (mGluR3) in dorsolateral prefrontal and entorhinal cortex, where they regulate cAMP– calcium opening of K⁺ channels to sustain neuronal firing and working memory. mGluR3 signaling is driven by N-acetylaspartylglutamate (NAAG) and constrained by glutamate carboxypeptidase II (GCPII), whose expression rises with age and inflammation. In prior work, chronic inhibition of GCPII with the orally bioavailable inhibitor 2-(3-mercaptopropyl) pentanedioic acid (2-MPPA) improved neuronal firing, working memory, and reduced pT217Tau pathology in aged macaques. Here, we employed liquid chromatography–tandem mass spectrometry (LC-MS/MS) to define the proteomic consequences of chronic 2-MPPA treatment in vulnerable (entorhinal cortex, dorsolateral prefrontal cortex) versus resilient (primary visual cortex) regions. We identified >2,400 proteins across experimental conditions, and label-free quantification revealed region-specific differential expression patterns paralleling known vulnerability gradients in sAD. Gene ontology enrichment of vulnerable regions implicated pathways governing protein deneddylation, amyloid and tau-associated processes, synaptic plasticity, mitochondrial homeostasis, and oxidative stress, revealing putative targets for therapeutic intervention in sAD. These findings demonstrate that GCPII inhibition engages distinct, region-selective molecular programs in the aging primate cortex, consistent with the protection of circuits most vulnerable to sAD. By mapping the proteomic shifts that occur with treatment, we reveal molecular signatures that not only serve as candidate biomarkers but also highlight novel mechanistic pathways contributing to calcium-driven degeneration in sAD. As such, more focused investigations into these pathways of therapeutic interest are warranted, in addition to the analysis of key post-translational modifications and their potential roles in sAD.

## Introduction

Improved therapeutic strategies are needed to reduce the risk and possibly prevent the neuropathology of sporadic Alzheimer’s disease (sAD). Although recent evidence suggests that anti-amyloid therapies can slow the progression of sAD, the findings remain quite subtle and additional approaches are needed to reduce toxic cascades at earlier stages of pathology prior to neuronal damage [1–3]. Dysregulated calcium signaling is well established as an early etiological factor in the pathogenesis of sAD and is linked to inflammatory mechanisms [4–6]. Thus, anti-inflammatory treatments aimed at restoring calcium regulation may help prevent or slow pathological cascades. However, interventions aimed at these upstream causes of sAD pathology should be tested in animal models with inflammatory and regulatory mechanisms similar to humans, and with naturally occurring calcium dysregulation and sAD-like pathology. Mouse models are inadequate for this purpose, as they use genetic modifications based on autosomal dominant disease that are downstream of calcium dysregulation in order to produce AD-like pathology. In contrast, macaques naturally develop cognitive deficits, calcium dysregulation, amyloid plaques, and tau pathology, including tangles in the oldest animals [7,8]. Importantly, mice have poorly developed association cortices, with molecular regulation and inflammatory actions that differ from primates in ways that may be central to the mechanisms that drive AD pathology.

Recently, our studies revealed an evolutionary expansion of type 3 metabotropic glutamate receptor (mGluR3)-*N*-acetylaspartylglutamate (NAAG)-glutamate carboxypeptidase II (GCPII) signaling in vulnerable sAD cortical circuits. mGluR3 are commonly found on astrocytes in both rodents and primates, where they potentiate glutamate uptake from the synapse. However, there are large species differences in mGluR3 location and function in neurons. In rodents, mGluR3 are concentrated on presynaptic receptors, where they reduce glutamate release [9,10]. However, in macaques, mGluR3 are predominately *postsynaptic* on dendritic spines in the dorsolateral prefrontal cortex (dlPFC) and entorhinal cortex (ERC) [10–12]. They have a critical, beneficial role at this location, where they regulate cAMP-calcium opening of K^+^ channels, and enhance, rather than reduce, neuronal firing and working memory performance [10,12]. Indeed, mGluR3 are often localized on the spine membrane near the smooth endoplasmic reticulum, positioned to regulate cAMP drive on internal calcium release. mGluR3s are stimulated not only by glutamate, but also by NAAG, which is co-released with glutamate and is selective for mGluR3 [13] (**Fig 1a**). However, under conditions of inflammation, microglia produce GCPII, which catabolizes NAAG and weakens mGluR3 regulation of calcium signaling [13–16] (**Fig. 1b**). This mechanism plays a large role in primates, as inhibition of GCPII in aged macaques with naturally-occurring inflammation markedly improved dlPFC neuronal firing, improved working memory, and with chronic treatment, reduced tau pathology in the dlPFC and ERC (**Fig 1c**). Specifically, we showed that daily treatment with the GCPII inhibitor 2-MPPA (2-[3-mercaptopropyl] pentanedioic acid) reduced levels of pT217Tau in the dlPFC and ERC and in the plasma of aged macaques, and that GCPII activity highly correlated with pT217Tau levels in the dlPFC. This data suggests that this mechanism, which expands in primates, significantly contributes to early-stage AD pathology. However, an understanding of the molecular substrates and underlying mechanisms of 2-MPPA efficacy is limited, especially at the proteomic level.

**Fig. 1.**
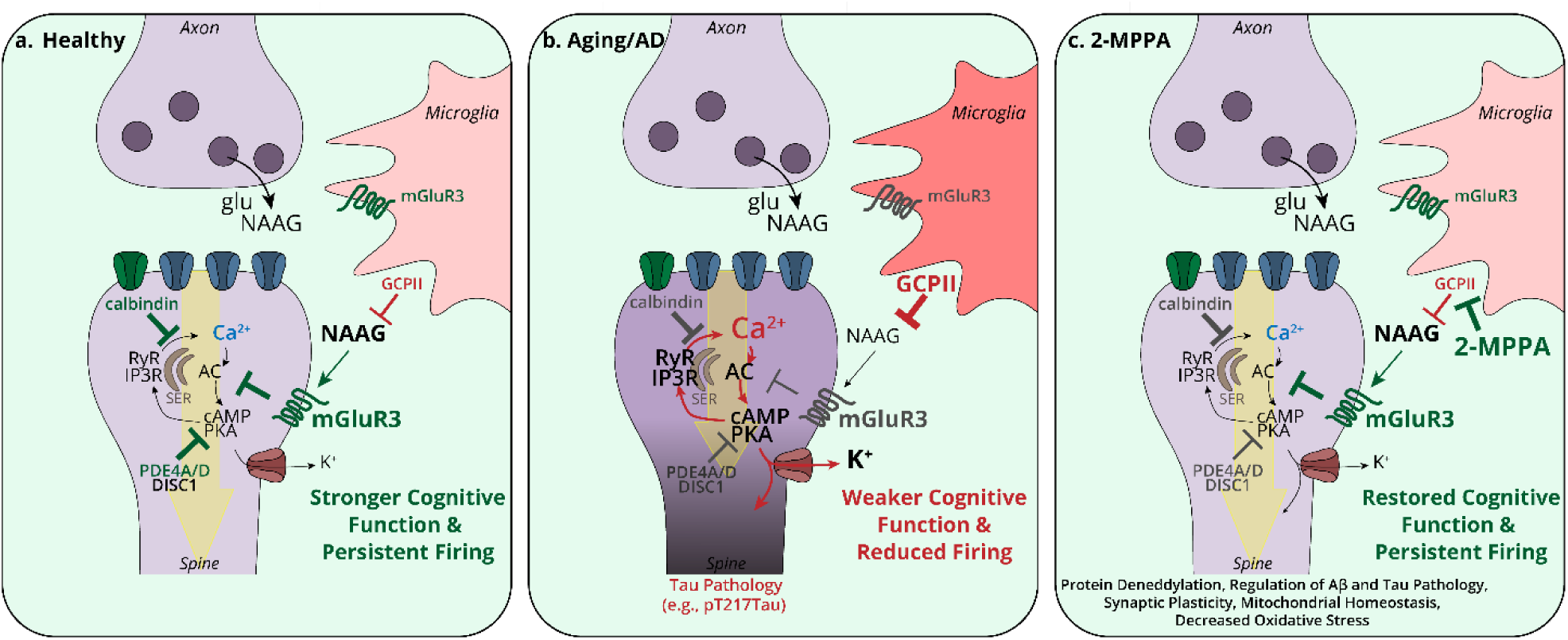
GCPII inhibition as a novel therapeutic strategy for sporadic Alzheimer’s disease. **a.** Schematic diagram highlighting how, under healthy conditions, postsynaptic compartments in association cortices (e.g., entorhinal cortex [ERC] and dorsolateral prefrontal cortex [dlPFC]) express the molecular machinery for feedforward cAMP–calcium signaling essential for network connectivity necessary for higher-order cognition. Feedforward calcium–cAMP signaling in ERC and dlPFC postsynaptic compartments is modulated by the phosphodiesterases (PDE4s), which catabolize cAMP, calcium buffering proteins such as calbindin, and by the Gi/o-linked receptors mGluR3 (shown) and α2A-AR (not shown), which are both predominantly localized on dendritic spines and inhibit the synthesis of cAMP. mGluR3 are stimulated by glutamate, but also by NAAG, which is co-released with glutamate and is specific for mGluR3. mGluR3 are also expressed in their traditional location on astrocytes (not shown), where they promote glutamate uptake. Microglia (light pink) also synthesize GCPII, an enzyme which catabolizes NAAG and reduces mGluR3 signaling. Under homeostatic healthy conditions, prominent NAAG stimulation of mGluR3 on dendritic spines, due to low levels of GCPII, enhances neuronal firing by inhibiting cAMP–PKA opening of K+ channels. In contrast, in classical glutamate synapses (e.g., primary visual cortex [V1]), neurotransmission depends on AMPA receptors, which provide permissive actions by depolarizing the postsynaptic membrane to eject Mg^2+^ from the NMDAR pore (green), thereby permitting NMDAR neurotransmission. The calcium entry through NMDAR can drive cAMP–PKA signaling to increase neuroplasticity and strengthen connections. **b.** Under conditions of inflammation/advancing age and in sAD, reactive microglia (darker pink) increase their expression of GCPII, which catabolizes NAAG and decreases mGluR3 regulation of cAMP–calcium signaling in ERC and dlPFC postsynaptic compartments. Aging/inflammation also reduce the expression of PDE4A/D and calbindin within postsynaptic compartments, resulting in raised levels of cAMP–PKA–calcium signaling, including PKA-mediated phosphorylation of ryanodine receptors to drive calcium “leak” from the SER. Excessive cAMP–PKA–calcium signaling opens K^+^ channels, which reduce neuronal firing, and high levels of cytosolic calcium activate calpain-2, which cleave and activate kinases such as GSK3β to hyperphosphorylate tau, for example, at pT217tau. **c.** 2-MPPA reduces GCPII expression to potentiate mGluR3-mediated regulation of calcium signaling within postsynaptic compartments to attenuate AD-related pathology. Our proteomic studies reveal that chronic 2-MPPA administration in aged rhesus macaques modulates pathways related to protein deneddylation, amyloid and tau pathology, synaptic plasticity, mitochondrial homeostasis, and oxidative stress, revealing putative targets for therapeutic intervention in sAD. cAMP, cyclic adenosine monophosphate; GCPII, glutamate-carboxypeptidase II; GSK3β, glycogen synthase kinase-3 beta; mGluR3, metabotropic glutamate receptor 3; NAAG, *N*-acetyl-aspartyl-glutamate; NMDAR, N-methyl-D-aspartate receptor; PKA, protein kinase A; p-tau, phosphorylated tau; SER, smooth endoplasmic reticulum.

In this study, we performed liquid chromatography-tandem mass spectrometry (LC-MS/MS) to evaluate proteomic signatures following chronic GCPII inhibition with 2-MPPA in vulnerable (ERC, dlPFC) and resilient (primary visual cortex (V1)) brain regions of sAD. Specifically, we leveraged archived brain tissue from the study described above wherein rhesus macaques received chronic 2-MPPA or placebo administration [17]. Proteomic studies in sAD have provided novel insights into disease pathways, discovery of biomarkers, and therapeutic targets [18–24]. Importantly, proteomics uniquely captures molecular signatures not detected with transcriptomic methods [19]. As such, we employed a bottom-up proteomics approach to identify protein content in each aforementioned brain region following treatment with 2-MPPA or placebo. In total, we identified over 2,400 proteins in at least three samples per condition. To better understand downstream effectors of 2-MPPA treatment, we used label-free quantification to identify differentially expressed proteins between conditions. Interestingly, differential expression mirrored known trends of region-specific susceptibility to AD pathogenesis. Finally, we employed gene ontology analysis to strengthen our understanding of the biological pathways implicated in 2-MPPA treatment. Overall, our findings reveal novel protein co-expression modules, thus allowing us to identify molecular substrates underlying GCPII inhibition in aging primate cortex.

## Materials and Methods

All research was approved by the Yale Institutional Animal Care and Use Committee (IACUC) and performed under National Institutes of Health (NIH) guidelines.

### Subjects

A total of 8 aged rhesus macaques (6 female, ages 20-28 yrs at start of study, 21-29 yrs at end) were used in the current study and were acquired from a Yale Cognitive Pharmacology lab that had retired. The animals were previously trained on a working memory task but had not been used in experimental studies for several years. Extremely old rhesus macaques, as used in the current study, are exceedingly rare, and thus the availability of these animals made this study possible.

### Drug treatment

2-MPPA (2-[3-mercaptopropyl]pentanedioic acid), is a brain-penetrant, orally-bioavailable thiol-based inhibitor of GCPII with an IC_50_ of 90nM which was first described in 2003 [25]. 2-MPPA’s pharmacokinetic profile has been detailed in both rodents [26] and humans [27].

In the current study, 2-MPPA (0.1 mg/kg, p.o.) was administered daily in a small piece of cereal bar treat; the technician administering the drug watched to ensure that the monkey ate all of the treat each day. Given that 2-MPPA can be unstable in solution, drug was diluted fresh each day in sterile saline immediately prior to administration [17]. On cognitive testing days (2 days per week), working memory testing and behavioral assessments were performed 2 hours after drug administration by a technician highly familiar with the normative behavior of each monkey but blind to drug treatment. The aged monkeys were tested on the delayed response task in a Wisconsin General Test Apparatus, a test of spatial working memory that is tightly associated with the integrity of the primate dorsolateral prefrontal cortex (dlPFC)[28–30]. Monkeys were treated with drug or vehicle on the day of sacrifice; the average time post-treatment to brain extraction was approximately 3 hrs (*vide infra*).

### Brain extraction

At the conclusion of the drug intervention paradigm, animals were euthanized by overdose of sodium pentobarbital (>150 mg/kg IV). Immediately after death, the scalp was incised and retracted to expose the cranium. The bone was cut in a circular manner starting at the superior level of the temporal bone. When necessary, depending on the thickness and hardness of the bone, a manual and/or electric autopsy saw was used to expediate bone removal. Sharp bony edges were removed with rongeurs or cut out with Littauer bone cutters. Once exposed, dura matter was lifted and removed by scissors. The brain was carefully exposed and immediately removed. The hemispheres were separated, and tissue blocks of about 1 cm in thickness were cut in the coronal plane with long tissue slide blades. Brain tissue blocks were immediately frozen.

### Mass spectrometry sample preparation

Frozen tissue was homogenized using the CryoCup Grinder (BioSpec Products, Cat. No. 206). Homogenized tissue was then transferred into ice-cold lysis buffer, consisting of 1% N-octylglucoside (Research Products International, N02207), 1% CHAPS (Research Products International, C41010), 0.5% sodium deoxycholate (Research Products International, D91500-25.0), 50 mM Tris, 100 mM NaCl, 2 mM MgCl_2_, benzonase (Sigma Aldrich, E1014), and protease inhibitor (Roche, 11836170001). Samples were rotated at 4 °C for 2 hrs, followed by centrifugation for 30 min at 15,000 rcf. Supernatants were collected and subjected to two sequential chloroform-methanol extractions to remove lipids and other contaminants. Briefly, a 4X sample volume of methanol (Fisher Chemical, A456-212) was added to supernatants, followed by 1X sample volume of chloroform (Avantor Sciences, 9180-01) and 3X sample volume of water (Fisher Chemical, Cat. No. W61), with thorough mixing after each addition. Samples were centrifuged for 1 min at 14,000 rcf, and the top aqueous layer decanted. Another 4X sample volume of methanol was added to the protein flake, vortexed, and centrifuged for 5 min at 20,000 rcf. Methanol was removed and protein pellet allowed to dry [31].

Following chloroform-methanol extraction, samples were resuspended in 150 µL of 50 mM ammonium bicarbonate (AmBic) (Honeywell Fluka, 40867). Samples were reduced with 1.5 mM dithiothreitol (DTT) (Sigma Aldrich, D0632) for 20 min at 65 °C, and then alkylated with 2.5 mM iodoacetamide (IAA) (Sigma Aldrich, I1149) in the dark for 15 min at RT. Subsequently, 1 µg of trypsin (Promega, V5111) was added and allowed to react overnight at 37 °C.

All reactions were quenched by adding 1 µL of formic acid (Thermo Scientific, 85178) and diluted to a volume of 200 µL prior to desalting. Desalting was performed using 10 mg Strata-X 33 µm polymeric reversed-phase SPE columns (Phenomenex, 8B-S100-AAK). Each column was activated with 500 µL of acetonitrile (ACN) (Honeywell, LC015) followed by 500 µL of 0.1% formic acid, 500 µL of 0.1% formic acid in 40% ACN, and equilibration with 500 µL of 0.1% formic acid. Samples were then added to the column and rinsed twice with 200 µL of 0.1% formic acid. Sample was eluted with 150 µL of 0.1% formic acid in 40% ACN, twice. Eluent was dried in a vacuum concentrator (LabConco) prior to reconstitution with 0.1% formic acid. Peptide concentrations were determined via NanoDrop One spectrophotometer (Thermo Fisher Scientific), and normalized amounts were prepared for mass spectrometry (MS) analysis.

### Mass spectrometry data acquisition

Samples were analyzed by online nanoflow liquid chromatography-tandem mass spectrometry using an Orbitrap Eclipse Tribrid mass spectrometer (Thermo Fisher Scientific) coupled to a Dionex UltiMate 3000 HPLC (Thermo Fisher Scientific). Reconstituted sample (2 µL) was injected onto an Acclaim PepMap 100 column packed with 2 cm of 5 µm C18 material (Thermo Fisher, 164564) with 0.1% formic acid in water (solvent A). Subsequently, peptides were separated on a 15 cm PepMap RSLC EASY-Spray C18 Column packed with 2 µm of C18 material (Thermo Fisher, ES904) using a gradient from 0 to 35% solvent B (0.1% formic acid, 80% ACN) in 60 min.

Full scan MS1 spectra were collected within a 400 to 1500 m/z mass range at a resolution of 60,000 and an automatic gain control (AGC) of 1e5. Dynamic exclusion was enabled with a repeat count of 2 and duration of 8 sec. Charge states 2 to 6 were selected for fragmentation. MS2s were generated at top speed for 3 sec. Higher-energy collisional dissociation (HCD) was performed on all selected precursor masses with an isolation window of 2 m/z, stepped collision energies of 25%, 30%, 40%, Orbitrap detection (resolution of 7500), maximum inject time of 75 msec, and a standard AGC target.

### Total protein analysis

Raw MS files were searched using Byonic against the full rhesus macaque proteome (Uniprot ID UP000006718). The mass tolerance was set to 10 ppm for MS1s and 20 ppm for MS2s. Met oxidation was set as a variable modification, and carbamidomethyl Cys was set as a fixed modification. Files were searched with fully specific cleavage C-terminal to Arg and Lys, with six allowed missed cleavages. A 1% protein FDR cutoff was applied. The mass spectrometry proteomics data was deposited to the ProteomeXchange Consortium via the PRIDE [32] partner repository with the data set identifier PXD065758.

Protein identifications from search outputs were compiled and reverse hits were removed. Results were filtered for a |LogProb| value greater than 1.3. For total protein numbers, identifications were considered true if they appeared in at least three samples per condition and/or region. (**Supp. Files 1-4** [ERC, dlPFC, V1, all regions, respectively])

### Label-free quantification

Label-free quantification was performed using the minimal workflow for MaxQuant according to the established protocol for standard data sets [33]. Files were searched against a focused database with fully-specific cleavage C-terminal to Arg and Lys, with 2 allowed missed cleavages. Carbamidomethyl Cys was set as a fixed modification. Raw data generated for each brain region was analyzed using Perseus according to the recommended protocol for label-free interaction data. Briefly, reverse and contaminant hits were removed from the data set and remaining values were log2 transformed. After grouping data according to condition, rows were filtered for valid values, i.e., those that represented more than one reported intensity. Missing values were imputed from normal distribution, and a two-sample T-test with an FDR >0.01 was applied. Proteins highlighted as significant correspond to a p-value < 0.05 and a |T-test Difference| > 1 (|fold change| > 2). (**Supp. File 5**)

### Gene ontology enrichment analysis

Protein lists generated from label-free quantification were sorted by values of “-Log p-value” multiplied by “Difference”. Positive and negative values, “upregulated” and “downregulated” with regard to 2-MPPA treatment respectively, were sorted by magnitude and input in rank order to geneontology.org. Analysis was performed against the “GO biological process complete” annotation data set for *Macaca mulatta*. The Bonferroni correction was applied for each analysis. Results were exported as a table and further analyzed in Microsoft Excel. (**Supp. File 6**)

### STRING analysis

Pathways of interest generated from gene ontology analysis were further visualized using string-db.org. Reference genes in designated pathways were input into STRING multiple protein viewer as their Uniprot ID. The full STRING network was visualized with line thickness indicating the strength of data support. A minimum required interaction score of 0.700 (high confidence) was applied. Query proteins only were visualized. Maps were exported as a vector graphic and further analyzed in Adobe Illustrator.

## Results

### Proteomic analyses of aging rhesus macaque brain regions following 2-MPPA or vehicle treatment

Aged rhesus macaques were treated with the GCPII inhibitor 2-MPPA daily for six months, as previously described (**Fig. 2a**). To identify protein changes that occurred in response to inhibition of GCPII, dissections of three brain regions, entorhinal cortex (ERC), dorsolateral prefrontal cortex (dlPFC), and primary visual cortex (V1), were processed for proteomic analysis. Each brain region was homogenized at liquid nitrogen temperatures, lysed, and subjected to chloroform-methanol protein extraction. Protein samples were digested with trypsin and analyzed via LC-MS/MS (**Fig. 2b**). In total, 2,428 proteins were identified in at least three samples per condition across brain regions, of which 2,022 proteins were identified in both treated and vehicle conditions. There were 250 unique proteins identified in tissues taken from 2-MPPA-treated macaques and 156 unique proteins in vehicle-treated tissue (**Fig. 2d**).

**Fig. 2.**
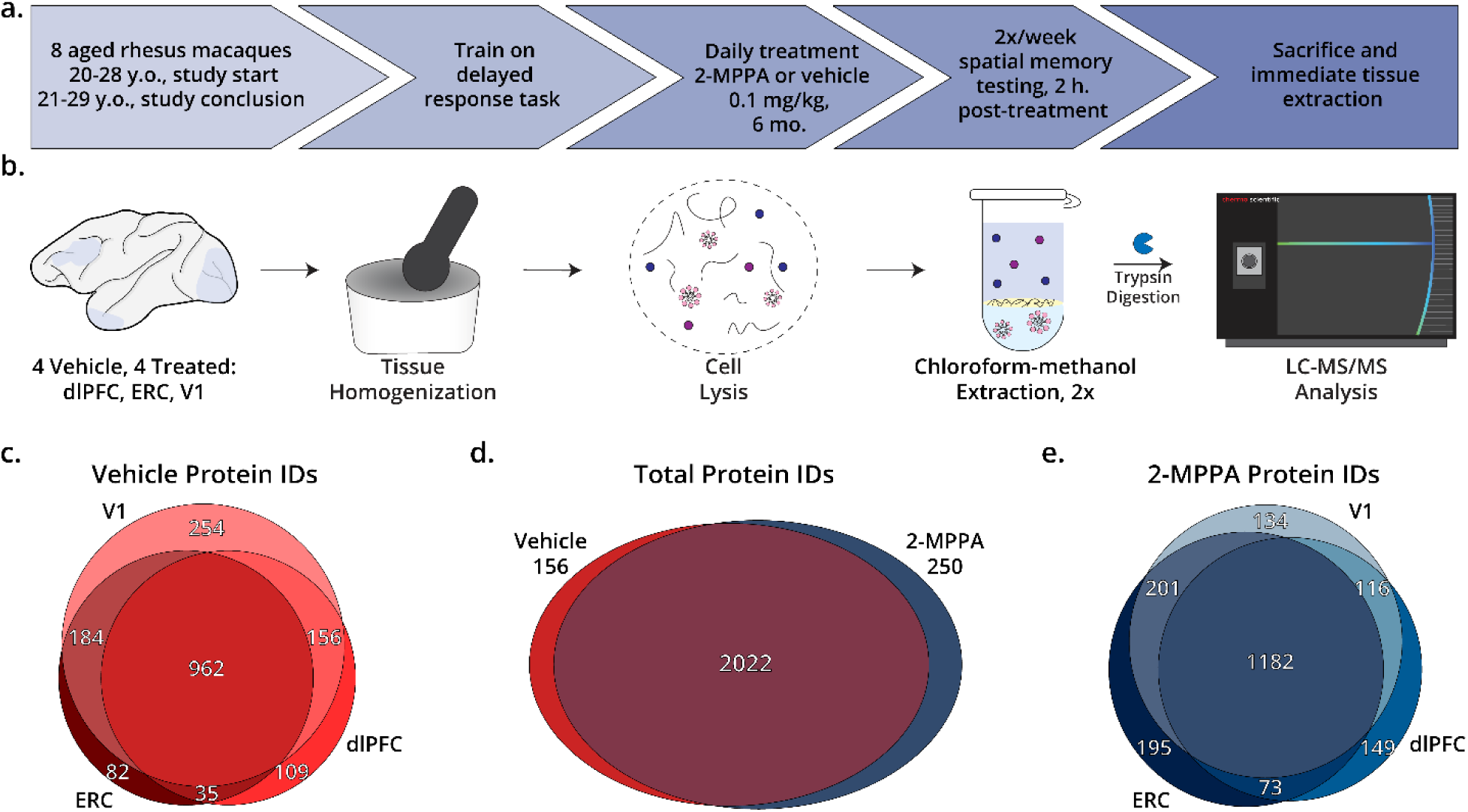
**a.** Treatment paradigm for rhesus macaques. **b.** Workflow for proteomics sample preparation. Frozen tissue blocks from dorsolateral prefrontal cortex (dlPFC), entorhinal cortex (ERC), and primary visual cortex (V1) were homogenized at liquid nitrogen temperatures and lysed. Tissue lysates were then subjected to two iterative chloroform-methanol extractions, digested with trypsin, and subjected to LC-MS/MS analysis. **c-e.** Raw MS files were searched in Byonic against the full rhesus macaque proteome. The mass tolerance was set to 10 ppm for MS1s and 20 ppm for MS2s. Met oxidation was set as a variable modification and carbamidomethyl Cys was set as a fixed modification. Files were searched with fully specific cleavage C-terminal to Arg and Lys, with 6 allowed missed cleavages. A 1% protein FDR cutoff was applied. Protein identifications were compiled and reverse hits removed. Results were filtered for p-value < 0.05. Protein identifications that appeared in at least three samples per region and/or condition were included in overall counts. Euler plots demonstrate counts and overlap of protein identifications between entorhinal cortex (ERC), dorsolateral prefrontal cortex (dlPFC), and primary visual cortex (V1) for both (**c**) vehicle-treated (n=4) and (**e**) 2-MPPA-treated (n=4) rhesus macaques. **d.** Euler plot demonstrating total protein identifications and overlap, regardless of region-specificity, between vehicle- and 2-MPPA-treated subjects.

In monkeys that received only placebo, a total of 1,782 proteins were identified in at least three samples per region. Of these, 82 unique proteins were identified in ERC, 109 in dlPFC, and 254 in V1. (**Fig. 2c**). In monkeys that received 2-MPPA treatment, 2,050 proteins were identified in at least three samples per region; 195 unique proteins were identified in ERC, 149 in dlPFC, and 134 in V1. (**Fig. 2e**). Interestingly, there was a greater deficit in protein identifications in the ERC (388) and dlPFC (258) of the vehicle-treated condition, compared to the V1 (77), reflective of established trends of region-specific vulnerability and pathological onset (**Supp. Fig. 1**) [34,35].

### Differential protein expression across vulnerable and resilient brain regions in sAD

To better understand the changes that occurred within each brain region in response to 2-MPPA treatment, we performed label-free quantification (LFQ), which allowed us to identify significantly altered protein levels. Proteins highlighted as significant reflect a fold-change greater than 2 and a p-value less than 0.05. Overall, we found the highest number of significantly changed proteins in ERC, followed by dlPFC, and finally V1. Again, this trend notably correlated with region-specific AD susceptibility and thus the trend of pathological onset (**Fig. 3a**) [34,35].

**Fig. 3.**
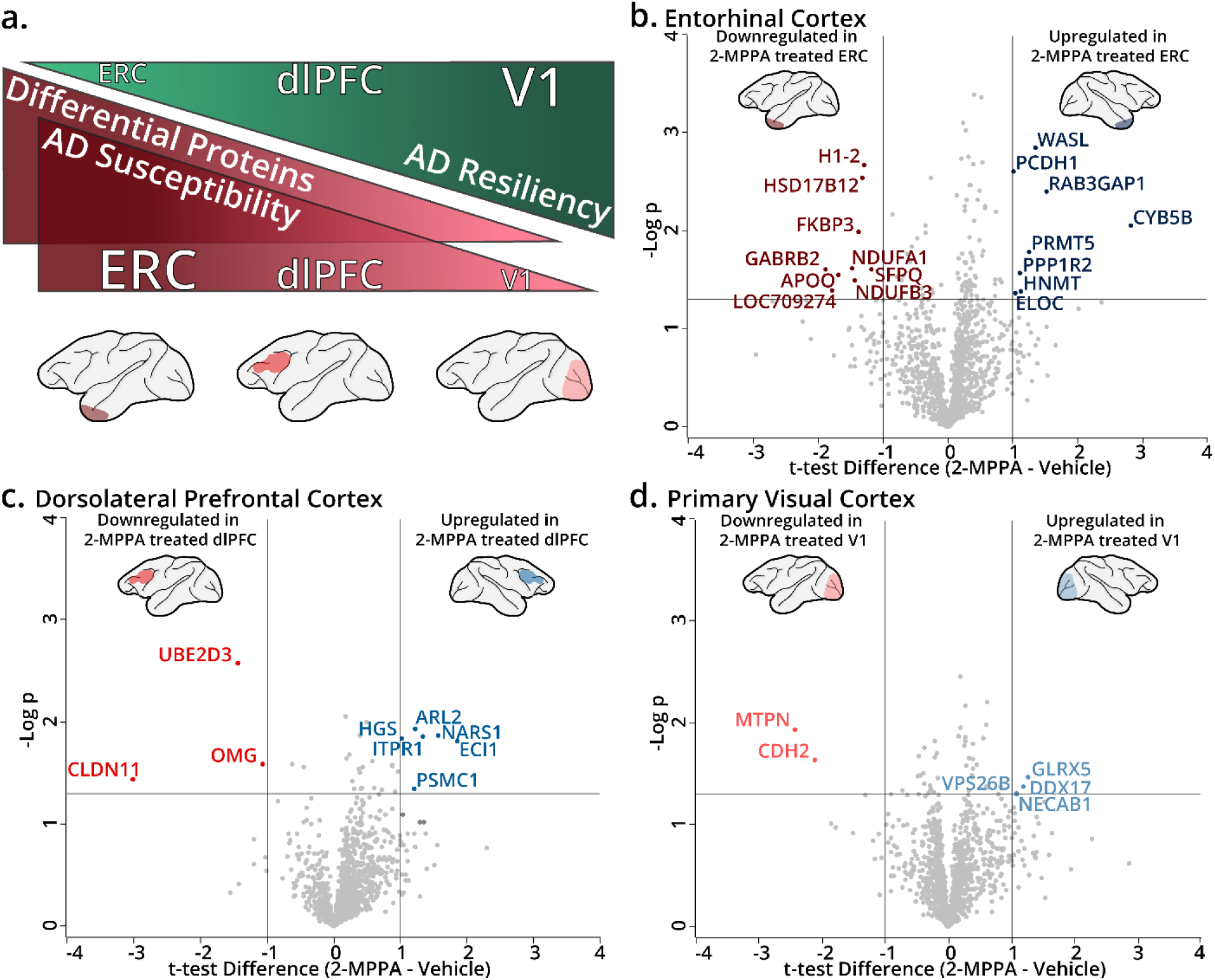
**a.** Schematic representation of region-specific AD susceptibility or resiliency, correlated to the number of differentially expressed proteins. Differential protein abundance for 2-MPPA-treated (n=4) versus vehicle-only control (n=4) observed in (**b**) entorhinal cortex, (**c**) dorsolateral prefrontal cortex, and (**d**) primary visual cortex. Label-free quantification was performed using the minimal workflow for MaxQuant according to the established protocol for standard data sets. Files were searched against a focused database with fully-specific cleavage C-terminal to Arg and Lys, with 2 allowed missed cleavages. Carbamidomethyl Cys was set as a fixed modification. Raw data generated for each brain region was analyzed using Perseus according to the recommended protocol for label-free interaction data. Briefly, reverse and contaminant hits were removed from the data set and remaining values were log2 transformed. After grouping data according to condition, rows were filtered for valid values, i.e., those that represented more than one reported intensity. Missing values were imputed from the normal distribution, and a two-sample T-test with an FDR > 0.01 was applied. Proteins highlighted as significant correspond to a p-value < 0.05 and a |T-test Difference| > 1 (i.e., |fold change| > 2).

#### Entorhinal cortex (ERC)

In ERC samples, nine proteins were found to be significantly downregulated following treatment with 2-MPPA (**Fig. 3b**). Of these, seven proteins were identified in existing AD proteomics literature, as reported in the NeuroPro database, including very-long-chain 3-oxoacyl-CoA reductase (HSD17B12), gamma-aminobutyric acid receptor subunit beta-2 (GABRB2), MICOS complex subunit (APOO), NADH dehydrogenase [ubiquinone] 1 beta subcomplex subunit 3 (NDUFB3), histone H1.2 (H1-2), peptidylprolyl cis-trans isomerase FKBP3 (FKBP3), and splicing factor proline- and glutamine-rich (SFPQ) [36]. Notably, three of these proteins are reported in the literature with opposite directionality to what we observed (**Table 1**) [36]. For example, histone H1.2 was consistently reported as upregulated in AD according to existing literature but was significantly downregulated in our 2-MPPA-treated cohort. Histone H1.2 is known to stabilize the aggregation of Aβ fibrils by binding to amyloid-like motifs and altering the conformation of beta-amyloid peptides, thus its downregulation could represent a promising downstream effect of 2-MPPA treatment [37]. Similarly, FKBP3, a peptidyl-prolyl *cis/trans* isomerase (PPIase), was uniquely downregulated in our 2-MPPA-treated dataset. PPIases represent a class of molecular chaperones that act by regulating protein folding at proline residues and modulating protein phosphorylation. Previous work has shown other proteins in this family, namely FKBP52 and FKBP51, promote tau oligomerization [38]. Finally, although conflicting evidence exists for the up- or downregulation of SFPQ in published proteomic datasets, there is substantial precedent for the dysregulation of this protein in AD and other tauopathies. In the context of AD, SFPQ mislocalizes from the nucleus to the cytosol of the cell, where it associates with tau oligomers in stress granules [39]. Our data suggests that SFPQ protein levels are downregulated in macaque ERC following 2-MPPA administration, suggesting that GCPII inhibition can attenuate tau pathology through multiple mechanisms.

**Table 1.** Comparison of differentially expressed ERC proteins to NeuroPro database. Up- and down-regulated proteins from ERC, relative to 2-MPPA treatment, identified by label-free quantification analysis that exceed threshold of p-value < 0.05 and |fold-change| > 2. Proteins identified by gene name. Bold text indicates proteins whose differential expression exists in contrast with established literature. Asterisk indicates literature study was performed in specific region of interest.

**Tables and Supplementary Figures**
| Gene Name | Direction of protein change in 2-MPPA cohort | Precedent in existing literature, relative to AD condition |
| --- | --- | --- |
| **H1-2** | Downregulated | Up (McKetney 2019* [106]; Xu 2019 [107]*; Wang 2020 [108]; Seyfried 2017 [109])<br>Present (Drummond 2022 [110], 2017 [111]) |
| HSD17B12 | Downregulated | Down (Johnson 2020 [112], 2022 [19]) |
| **FKBP3** | Downregulated | Up (Hondius 2016 [113])<br>Present (Drummond 2022 [110], 2017 [111]) |
| NDUFA1 | Downregulated | N/A |
| GABRB2 | Downregulated | Down (Johnson 2020 [112], 2022 [19]; Wingo 2020 [114]) |
| **SFPQ** | Downregulated | Down (Astillero-Lopez 2022* [115]; Seyfried 2017 [109])<br>Up (Johnson 2020 [112])<br>Present (Drummond 2017 [111], 2020 [116], 2022 [110]) |
| APOO | Downregulated | Down (Seyfried 2017 [109]; Johnson 2020 [112], 2022 [19]; Andreev 2012 [117]; Drummond 2022 [110])<br>Present (Drummond 2017 [111]) |
| NDUFB3 | Downregulated | Down (Carlyle 2021 [90]; Johnson 2022 [19]; Seyfried 2017 [109])<br>Present (Drummond 2017 [111], 2022 [110]) |
| LOC709274 | Downregulated | N/A |
| WASL | Upregulated | Up (Xu 2019 [107])<br>Present (Drummond 2017 [111], 2022 [110]) |
| **PCDH1** | Upregulated | Down (Hondius 2016 [113]; Johnson 2018 [118], 2020 [112], 2022 [19]; Mendonca 2019 [119]; Seyfried 2017 [109])<br>Present (Drummond 2017 [111], 2022 [110]) |
| **RAB3GAP1** | Upregulated | Down (Mendonca 2019* [119], Johnson 2018 [118], 2022 [19]) |
| CYB5B | Upregulated | Present (Drummond 2022 [110]) |
| PRMT5 | Upregulated | Up (Xu 2019 [107]; Mendonca 2019 [119])<br>Present (Drummond 2022 [110]) |
| **PPP1R2** | Upregulated | Down (Mendonca 2019 [119]; Johnson 2020 [112]) |
| HNMT | Upregulated | Up (McKetney 2019* [106]; Johnson 2022 [19]; Xu 2019 [107]) |
| **ELOC** | Upregulated | Down (Johnson 2022 [19]; Xu 2019 [107]) |

\*Specified region of interest

Eight proteins in ERC were significantly upregulated in our dataset following 2-MPPA treatment: actin nucleation-promoting factor WASL (i.e., neural Wiskott-Aldrich syndrome protein), protocadherin-1 (PCDH1), Rab3 GTPase-activating protein catalytic subunit (RAB3GAP1), cytochrome b5 type B (CYB5B), protein arginine N-methyl transferase 5 (PRMT5), protein phosphatase inhibitor 2 (PPP1R2), histamine N-methyl transferase (HNMT), and elongin C (ELOC) (**Figure 3b**). Each of these proteins was observed in other proteomic analyses (**Table 1**). RAB3GAP1 showed opposite directionality to ours in three studies, i.e., upregulated in our study but downregulated in others [36]. Rab3 GTPase-activating proteins have been shown to play key roles in neurotransmitter release and synaptic plasticity, the latter of which is a key deficiency in AD. As such, its upregulation in our dataset could imply a promising shift towards enhancing neuronal signaling [40]. Similarly, the upregulation of PCDH1 suggests 2-MPPA may be strengthening neuronal connections via upregulation of protocadherins, though we note only one of these was detected in our study [41–44]. Other upregulated proteins also demonstrated potential protective effects of 2-MPPA treatment. For example, WASL is involved in myelin wrapping of oligodendrocytes; demyelination is a common feature of neurodegenerative diseases, including AD [45]. WASL also modulates actin cytoskeleton polymerization in conjunction with cell division cycle 42 (CDC42) by stimulating the actin-nucleating activity of the actin-related protein 2/3 (ARP2/3) complex to regulate dendritic spine morphogenesis and plasticity [46–49]. These findings suggest that 2-MPPA might promote synaptic plasticity by enhancing actin cytoskeleton dynamics. Finally, PRMT5 protects against neuronal cell death and is downregulated in the presence of Aβ [50]. Together, these results indicate potential therapeutic effects of 2-MPPA treatment through both the downregulation of Aβ and tau stabilizing interactors and the upregulation of various mediators of synaptic function.

#### Dorsolateral prefrontal cortex (dlPFC)

In dlPFC, the expression of nine proteins was significantly altered between conditions (**Fig. 3c**). Three proteins, E2 ubiquitin-conjugating enzyme (UBE2D3), claudin (CLDN11), and oligodendrocyte-myelin glycoprotein (OMG), were found to be downregulated in 2-MPPA treated samples. Conflicting results exist for these proteins in other AD proteomic studies (**Table 2**) [36]. However, their downregulation with treatment in our dataset is a promising result. Namely, CLDN11 has been shown to significantly increase in AD frontal cortex neurons [51].

**Table 2.** Comparison of differentially expressed dlPFC proteins to NeuroPro database. Up- and down-regulated proteins from dlPFC, relative to 2-MPPA treatment, identified by label-free quantification analysis that exceed threshold of p-value < 0.05 and |fold-change| > 2. Proteins identified by gene name. Bold text indicates proteins whose differential expression exists in contrast with established literature. Asterisk indicates literature study was performed in specific region of interest.

| Gene Name | Direction of protein change in 2-MPPA cohort | Precedent in existing literature, relative to AD condition |
| --- | --- | --- |
| **UBE2D3** | Downregulated | Down (Johnson 2020* [112], 2022* [19]; Mendonca 2019 [119])<br>Up (Zellner 2022 [120]) |
| **CLDN11** | Downregulated | Down (Hales 2016* [121]; Johnson 2020 [112], 2022 [19])<br>Up (Johnson 2018* [118])<br>Present (Drummond 2022 [110]) |
| **OMG** | Downregulated | Down (Li 2021 [122]; Zhang 2018* [24])<br>Up (Johnson 2018* [118])<br>Present (Drummond 2017 [111], 2022 [110]) |
| ARL2 | Upregulated | Present (Drummond 2022 [110]) |
| **NARS1** | Upregulated | Down (Johnson 2020* [112], 2022* [19])<br>Up (Zellner 2022 [120])<br>Present (Drummond 2017 [111], 2022 [110]) |
| **ITPR1** | Upregulated | Down (Johnson 2022* [19]; Xu 2019 [107])<br>Up (Ho Kim 2015 [123])<br>Present (Drummond 2022 [110]) |
| **HGS** | Upregulated | Down (Johnson 2018* [118], 2020* [112], 2022* [19]; Seyfried 2017* [109]; Xu 2019 [107])<br>Up (Zellner 2022 [120])<br>Present (Drummond 2022 [110]) |
| **ECI1** | Upregulated | Down (Johnson 2022* [19])<br>Up (Xu 2019 [107])<br>Present (Drummond 2022 [110]) |
| PSMC1 | Upregulated | Present (Drummond 2017 [111], 2022 [110]) |

\*Specified region of interest

Conversely, six proteins in dlPFC were found to be upregulated following treatment: ADP-ribosylation factor-like protein 2 (ARL2), asparagine-tRNA ligase (NARS1), inositol 1,4,5-triphosphate receptor (ITPR1), hepatocyte growth factor-regulated tyrosine kinase substrate (HGS), enoyl-CoA delta isomerase 1 (ECI1), and 26S protease regulatory subunit 4 (PSMC1). The differential expression of PSMC1 was particularly encouraging, as it is a subunit of the 26S proteasome necessary for degrading misfolded and damaged proteins, and its depletion has been linked to neurodegeneration and Lewy body formation [52]. Within the frontal cortex specifically, NARS1, ITPR1, HGS, and ECI1 have been observed as downregulated in AD, in contrast to our 2-MPPA treated macaques where levels were upregulated [36]. Interestingly, ITPR1, an intracellular calcium channel, is among a family of proteins that play key roles in modulating calcium signaling [53,54]. In fact, for decades, aberrant ITPR function has been detected in AD samples and Aβ accumulation was directly linked to its dysfunction [55–57]. While this change may challenge our hypotheses on the direct modes of action of GCPII in attenuating intracellular calcium dysfunction, the upregulation of ITPR1 might represent compensatory alterations, or might reflect changes in specific cell-types that are not possible to parse out with bulk proteomics. Furthermore, ECI1 is necessary for endothelial cell migration, and thus is tied to the formation/maintenance of microvasculature and the blood brain barrier (BBB). As such, ECI1 dysregulation in sAD might contribute to microvascular changes and BBB disruption [58–60]. Taken together, each of these potentially restored functions suggests a homeostatic shift following 2-MPPA treatment, further supporting potential neuroprotective effects as seen in the ERC.

#### Primary visual cortex (V1)

Finally, only six proteins were identified as significantly altered in V1, two of which were downregulated and four upregulated in 2-MPPA treated animals (**Fig. 3d**). Notably, all of these proteins were opposite to expression changes in existing AD proteomics studies (**Table 3**) [36]. For example, cadherin-2 (CDH2) is reported as present and/or upregulated in AD brains [36]. It has been shown that the deposition of Aβ plaques decreases the surface level of CDH2, increasing degradation products, and it has been suggested that neural function is disrupted partially through the interference of CDH2-mediated synaptic plasticity [61,62]. On the other hand, VPS26 retromer complex component B (VPS26B) is downregulated in two AD proteomics datasets [36]. In fact, the deficiency of VPS26B has been associated with region-specific synaptic vulnerability, as its depletion in the trans-entorhinal cortex is highly correlated with sortilin-like receptor 1 (SORL1) deficiency [63]. Mutations in SORL1 are strong genetic risk factors for AD [64].

**Table 3.** Comparison of differentially expressed V1 proteins to NeuroPro database. Up- and down-regulated proteins from V1, relative to 2-MPPA treatment, identified by label-free quantification analysis that exceed threshold of p-value < 0.05 and |fold-change| > 2. Proteins identified by gene name. Bold text indicates proteins whose differential expression exists in contrast with established literature. Asterisk indicates literature study was performed in specific region of interest.

| Gene Name | Direction of protein change in 2-MPPA cohort | Precedent in existing literature, relative to AD condition |
| --- | --- | --- |
| **MTPN** | Downregulated | Up (Andreev 2012 [117])<br>Present (Drummond 2022 [110]) |
| **CDH2** | Downregulated | Up (Hales 2016 [121])<br>Present (Drummond 2022 [110]) |
| **GLRX5** | Upregulated | Down (Mendonca 2019 [119]; Xu 2019* [107]) |
| **DDX17** | Upregulated | Down (Mendonca 2019 [119])<br>Up (Johnson 2020 [112], 2022 [19])<br>Present (Drummond 2020 [116], 2022 [110]) |
| **VPS26B** | Upregulated | Down (Andreev 2012 [117]; Mendonca 2019 [119])<br>Present (Drummond 2017 [111], 2022 [110]) |
| **NECAB1** | Upregulated | Down (Johnson 2020 [112], 2022 [19]; Xu 2019* [107])<br>Up (Zellner 2022 [120]) |

\*Specified region of interest

As mentioned, the lower number of differentially expressed proteins is indicative of a well-established trend within the AD brain wherein certain regions, such as the primary visual cortex, are more resilient to disease pathology than others [34,35]. This is reflected by the onset of AD symptoms, specifically the appearance of memory and cognitive deficits over time [65,66].

### Pathway Analysis

With LFQ data in-hand, we performed gene ontology (GO) analysis to more comprehensively understand changes that may be occurring in 2-MPPA treated macaques [67,68]. For this, we considered both the difference in expression (T-test difference/log_2_ fold change) and the significance (p-value) of the differences. Rank ordered lists based on these two factors were used for analysis of up- and downregulated proteins in each brain region relative to 2-MPPA treatment.

Given the established vulnerability and observed responsiveness of the ERC, we were particularly interested in the pathway enrichment analysis of this region. Thus, the top 20 up- and downregulated biological processes following 2-MPPA treatment can be seen in **Fig 4a and 4b**. Upregulated processes of note included protein de-neddylation, regulation of amyloid fibril formation, negative regulation of Aβ formation, negative regulation of amyloid precursor protein (APP) catabolic process, and synaptic vesicle localization. Protein neddylation is a ubiquitin-like modification involved in regulating various biological processes, particularly related to DNA replication, cell-cycle re-entry, and cell death [69]. In the context of AD, it is hypothesized that NEDD8, a ubiquitin-like protein, translocates from the nucleus to the cytoplasm [70]. Here, amyloid precursor protein binding protein-1 (APP-BP1) activates neddylation, contributing to the re-entry of neurons back into the cell cycle, leading to autophagic degeneration and neuronal cell death [71–74]. By enhancing protein deneddylation, 2-MPPA may serve to protect against neurodegeneration by acting on this pathway, specifically by attenuating the aberrant re-entry of neurons into the cell cycle. Similarly, enriched processes involving the regulation of Aβ formation and oligomerization are promising indications that 2-MPPA is ameliorating AD pathology. This is further exemplified by indications of synaptic function (“synaptic vesicle lumen acidification”, “synaptic vesicle localization”), which could suggest a recovery of normal synaptic activity and transmission following drug treatment. As synaptic dysfunction and neuronal atrophy are central to AD pathology, and correlate with cognitive deficits, the potential recovery of this network following 2-MPPA treatment is an encouraging therapeutic effect [75–78]. Conversely, downregulated pathways in the ERC suggested various mitochondrial and metabolic changes that occur in response to 2-MPPA treatment. Of note, the decrease in proteins associated with oxidative phosphorylation processes and cell redox homeostasis may indicate that 2-MPPA relieved the oxidative stress characteristic of AD/aging brains [79–82]. Similarly, we identified a downregulation in various components of the mitochondrial electron transport chain (ETC) in macaque ERC, such as “NADH to ubiquinone”, “ubiquinol to cytochrome c”, “mitochondrial and ATP synthesis coupled electron transport”, “aerobic ETC”, and “mitochondrial respiratory chain complex I assembly” (**Fig. 4b**). Downregulation of the ETC pathways following GCPII inhibition with 2-MPPA could reduce neuronal metabolic demand and reactive oxygen species production, promoting a shift toward a more energy-efficient, stress-resistant state. At the circuit level, this may help preserve synaptic integrity and functional connectivity under conditions of metabolic or oxidative stress, thereby stabilizing network activity and resilience in vulnerable brain regions [83,84].

**Fig. 4.**
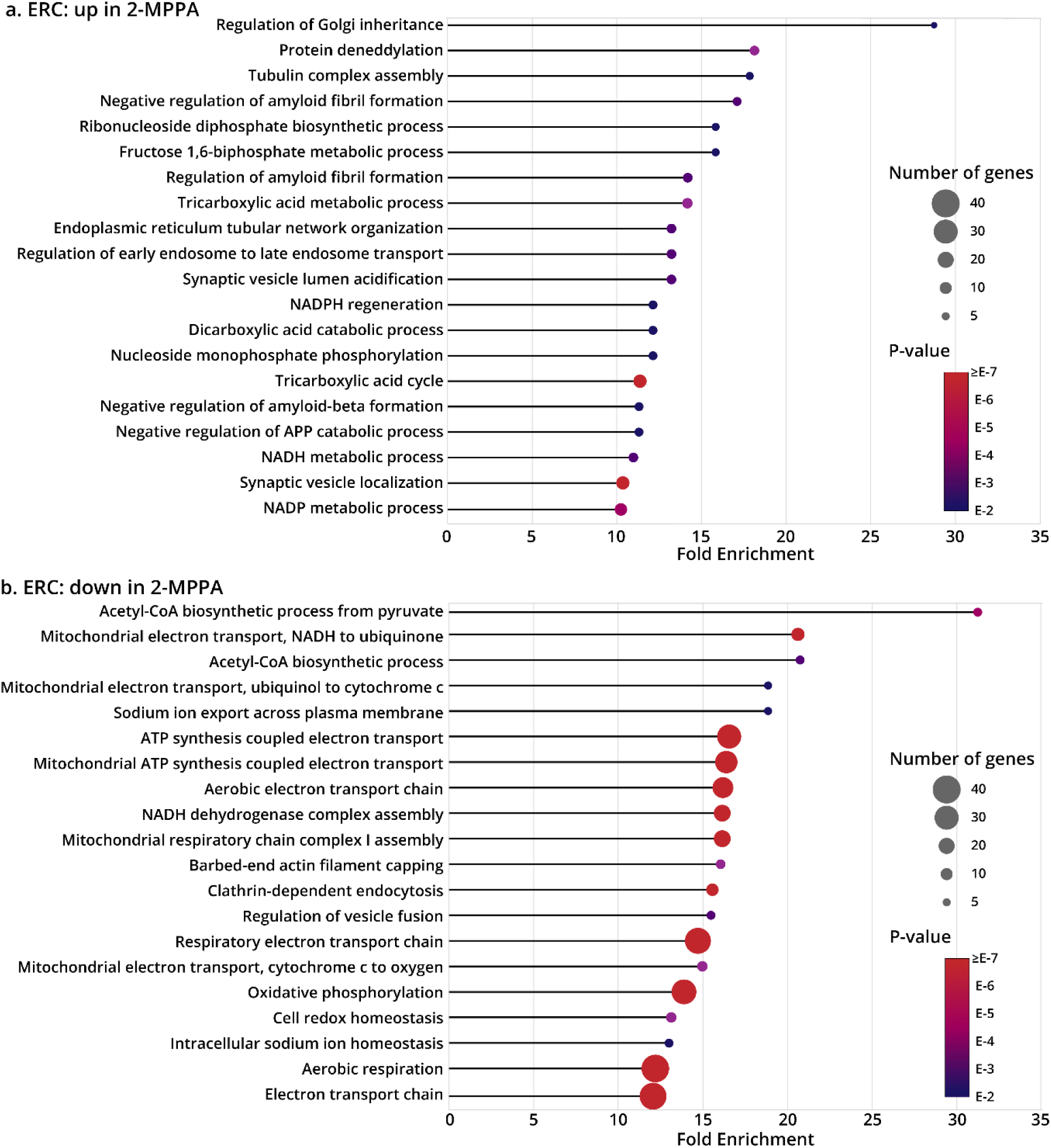
Gene ontology analysis of ERC proteins (**a**) upregulated or (**b**) downregulated with 2-MPPA treatment. Input protein lists, generated from label-free quantification analysis, were sorted according to both significance (p-value) and magnitude of difference (fold-change). Rank order proteins lists were analyzed using GO enrichment analysis for biological processes in *Macaca mulatta*, with the Bonferroni correction applied. The top 20 hits in either direction are represented.

Pathway enrichment analysis was similarly performed on both dlPFC (**Supp. Fig. 1**) and V1 (**Supp. Fig. 2**) datasets. Upregulated pathways of interest in dlPFC included a number of processes associated with synaptic function, such as synaptic vesicle recycling, transport, endocytosis, and localization (**Fig. 5**). The prevalence of DNA-related pathways (protein localization to telomere, regulation of protein localization to chromosome, telomere repair, etc.) downregulated in dlPFC with 2-MPPA treatment are similarly promising (**Supp. Fig. 2b**). As mentioned above, the aberrant re-entry of neurons into the cell cycle is a feature of AD, contributing to neuronal cell death as observed in rodent models of AD [73,74]. The downregulation of processes associated with DNA synthesis and replication could therefore suggest an alleviation of this phenotype. As demonstrated in the previous section, protein changes in V1 were markedly less than in the other brain regions. As such, GO analysis between conditions lacked the statistical power to truly identify significantly altered pathways. However, certain identifications could lend themselves towards representing the inherent resiliency of V1, such as the negative regulation of Aβ formation and APP catabolism (**Supp. Fig. 3a**).

**Fig. 5.**
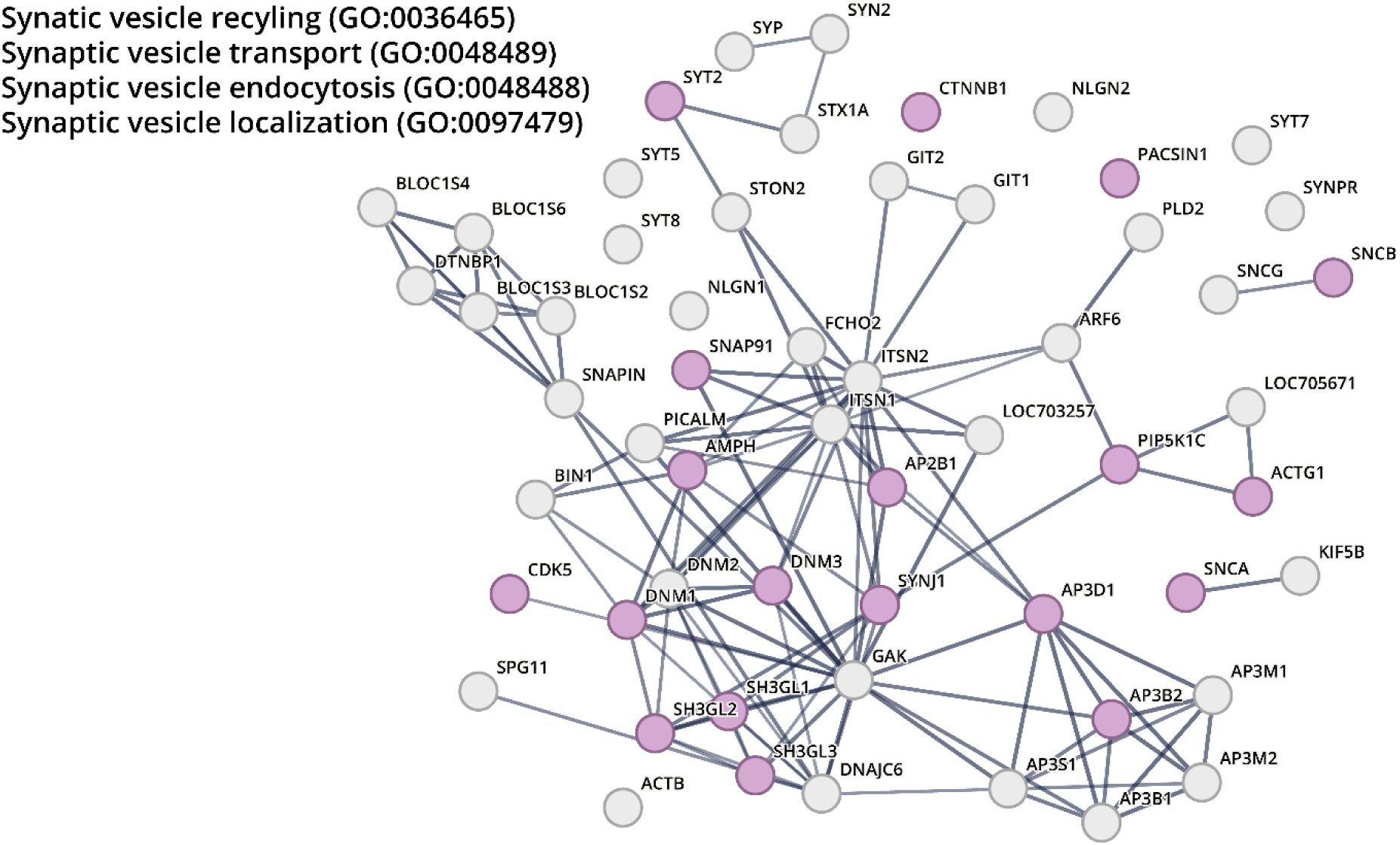
STRING protein network analysis for upregulated GO pathways of interest in 2-MPPA-treated dlPFC tissue. Reference genes from four pathways (*top left*) were queried and visualized, with line thickness indicating strength of data support. A minimum required interaction score of 0.700 (high confidence) was applied. Genes identified in our dataset are indicated with purple shading.

Overall, our results suggested that 2-MPPA treatment could serve to ameliorate AD pathology through various neuroprotective mechanisms, particularly with regard to synaptic maintenance and Aβ toxicity. While 6 months treatment of aged rhesus macaques with 2-MPPA resulted in modest changes, those that were observed show promise for the use of this drug as an early-stage therapeutic to hinder disease progression.

## Discussion

In this study, we provided proteomic analysis of aged rhesus macaques following chronic administration with a GCPII inhibitor, 2-MPPA, compared to vehicle-treated animals. Our results identified several signaling pathways and relevant proteins underlying the potential therapeutic efficacy of GCPII inhibition in augmenting cognition in higher-order cortical circuits. Our findings were consistent with the spatial and temporal progression of pathology in sAD, revealing most alterations in the highly vulnerable ERC, modest changes in dlPFC, and subtle alterations in the exceedingly resilient V1. Several of the identified proteomic changes following 2-MPPA treatment are particularly relevant for the pathogenesis of sAD and have been implicated in previous studies, as detailed above. Finally, biological pathway analysis highlighted a number of promising potential downstream effects of 2-MPPA treatment, for example, with regard to tau pathogenesis, APP/amyloid processing, and synaptic function (**Fig. 1c**). These findings complement the overarching hypothesis that anti-inflammatory GCPII inhibition helps to reduce AD pathology by restoring calcium regulation and lend additional credence to GCPII inhibition as a potential therapeutic strategy for sAD.

Proteomic studies in sAD have provided novel insights into disease pathways, discovery of biomarkers, putative therapeutic targets, disease staging, and etiology [18–24]. In fact, proteomics has revealed modules/pathways that are not observed using transcriptomics and snRNA-seq analyses in the same cohorts and brain regions, highlighting the unique contributions of proteomics in identifying molecular substrates underlying sAD pathogenesis. The proteomics dataset generated from the current study provides insights into how 2-MPPA might ameliorate perturbations that mediate neuropathology in sAD. For example, the upregulation of protein deneddylation could provide a mechanism whereby GCPII inhibition with chronic 2-MPPA administration can attenuate amyloid pathology through ubiquitin-like modifications, as detailed above [69–74]. Furthermore, the downregulation of oxidative phosphorylation and cell redox homeostasis might be significant in allaying the development and progression of sAD. Various studies have revealed that oxidative stress can accelerate the formation of amyloid plaques and NFTs in AD by abrogating mitochondrial function, leading to decreased energy metabolism and the generation of reactive oxygen species (ROS) that lead to a vicious cycle to further exacerbate oxidative stress pathways [80–82].

Synapse loss is well-defined, early, and robust pathology in AD [85–87]. It is considered to be a key structural correlate of cognitive deficits in the illness [75–78,88]. Histopathological examination of postmortem tissue and brain biopsies have revealed a decrement in synapse density in numerous cortical regions, including the hippocampus, PFC, ERC, cingulate gyrus, and temporal cortex in AD compared to control subjects [89]. This anatomical pattern has been observed in both the prodromal stage and in advanced patients with dementia. Proteomics studies have further identified various modules revealing synaptic dysfunction in aging and AD [90–92]. As a result, this has spurred numerous clinical trials to rescue synaptic plasticity deficits in AD [93,94]. Our proteomics findings suggest that chronic administration of 2-MPPA might enhance synaptic plasticity pathways, particularly in the newly evolved primate dlPFC. This region contains microcircuits, especially in deep layer III, with extensive recurrent excitation that generate the persistent neuronal firing needed to maintain mental representations in working memory.

Alterations in mitochondrial ETC function, like we identified with GO analysis following treatment, have complex and context-dependent consequences for neuronal viability. Potent and *chronic* inhibition of ETC components, such as with rotenone or MPP⁺, reliably produces dopaminergic neuron loss and α-synuclein aggregation, recapitulating Parkinson-like pathology in animal models rather than conferring neuroprotection [95,96]. These studies underscore that sustained ETC blockades are neurotoxic and primarily used to model disease mechanisms. In contrast, emerging evidence suggests that *partial* or transient modulation of ETC activity may have beneficial effects by engaging adaptive stress pathways. For example, mild inhibition of Complex I can activate AMPK signaling, promote mitophagy, augment antioxidant activity, and restore bioenergetic balance— mechanisms implicated in the actions of metformin and related compounds. In symptomatic APP/PS1 mice, partial Complex I inhibition improved energy homeostasis, synaptic function, and cognition while reducing oxidative stress and neuroinflammation [83]. Consistently, metformin use in humans has been associated with reduced risk of cognitive impairment and dementia [97], and preclinical studies demonstrate that metformin ameliorates Aβ-induced cognitive deficits, attenuates oxidative and inflammatory stress, and enhances autophagic activity [84,98]. Together, these findings indicate that while chronic ETC inhibition is deleterious, strategic modulation of mitochondrial metabolism with 2-MPPA—potentially analogous to metformin’s mild Complex I inhibition—could engage protective signaling pathways relevant to neurodegenerative disease mechanisms.

Given our own analyses identified a number of biologically-significant pathways affected by 2-MPPA administration, it would be useful to perform more focused investigations into these effects. Namely, the effect of 2-MPPA treatment on synaptic function is of particular therapeutic relevance. Additionally, a more in-depth analysis of key post-translational modifications (PTMs), such as phosphorylation and glycosylation, could yield interesting findings into the role of these modifications in sAD. Finally, as GCPII is especially active in inflammatory states, we hypothesize 2-MPPA inhibition will induce changes to this overall state in aging brains. Studies of inflammatory signatures in the blood prior to and following chronic 2-MPPA administration could identify novel inflammatory biomarkers for earlier detection of sAD.

As discussed above, rhesus macaques offer distinct advantages for studying the etiology of sAD, particularly due to their genetic and anatomical similarities to humans, and for investigating potential disease-modifying therapies that are upstream of canonical pathological hallmarks [99–102]. Although rhesus macaques do not develop the same magnitude of fibrillated tau pathology as humans, they do enable the study of early-stage, soluble tau pathology, such as pT217Tau, which has emerged as a novel fluid-based biomarker and heralds future neurodegeneration [103,104]. In fact, our previous studies revealed that chronic 2-MPPA administration robustly decreased pT217Tau levels in vulnerable brain regions and in blood plasma, highlighting efficacy in ameliorating pathological tau species relevant to the etiology of sAD. As such, further studies leveraging this model system are warranted.

That said, it is pertinent to note a critical limitation of the current study. As mentioned, aged rhesus macaques are exceedingly rare, and thus we were notably hampered by the number of subjects we could investigate. To mitigate this, we applied a more stringent fold-change cutoff compared to other AD proteomic studies, which investigate sample sizes sometimes into the hundreds. As such, more nuanced changes, such as those that reach T-test statistical significance but not difference thresholds, were not highlighted by our study. However, the overall trends we presented represent a promising shift towards homeostasis in treated monkeys.

Overall, our study presents a unique paradigm for investigating therapeutic interventions upstream of typical pathological hallmarks. The beneficial effects of GCPII inhibitors appear to be an evolutionary advance, as mGluR3-NAAG-GCPII signaling is associated with higher cognitive function in humans.[105] The use of mass spectrometry-based proteomics enabled the identification of a number of biologically-relevant proteins and pathways modulated by chronic GCPII inhibition, thus supporting the use of GCPII inhibitors in early intervention of sAD.

## Supplementary Figures

**Supp. Fig. 1.**
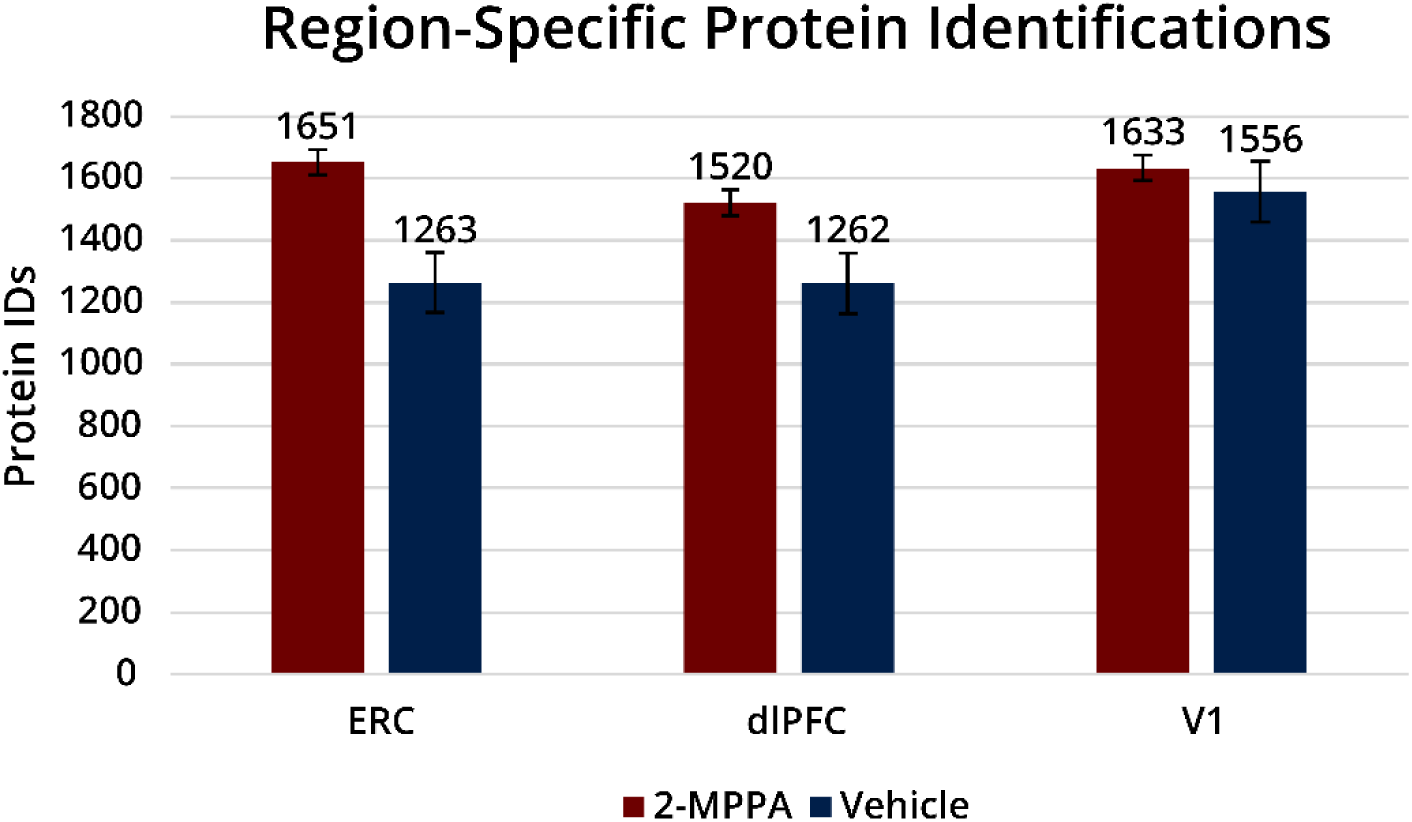
Total protein identifications for each brain region, per condition. Raw MS files were searched in Byonic against the full rhesus macaque proteome. The mass tolerance was set to 10 ppm for MS1s and 20 ppm for MS2s. Met oxidation was set as a variable modification and carbamidomethyl Cys was set as a fixed modification. Files were searched with fully specific cleavage C-terminal to Arg and Lys, with 6 allowed missed cleavages. A 1% protein FDR cutoff was applied. Protein identifications were compiled and reverse hits removed. Results were filtered for p-value < 0.05. Protein identifications that appeared in at least three samples per region and condition were included in counts. Bar graphs were generated in Excel with standard error bars included.

**Supp. Fig. 2.**
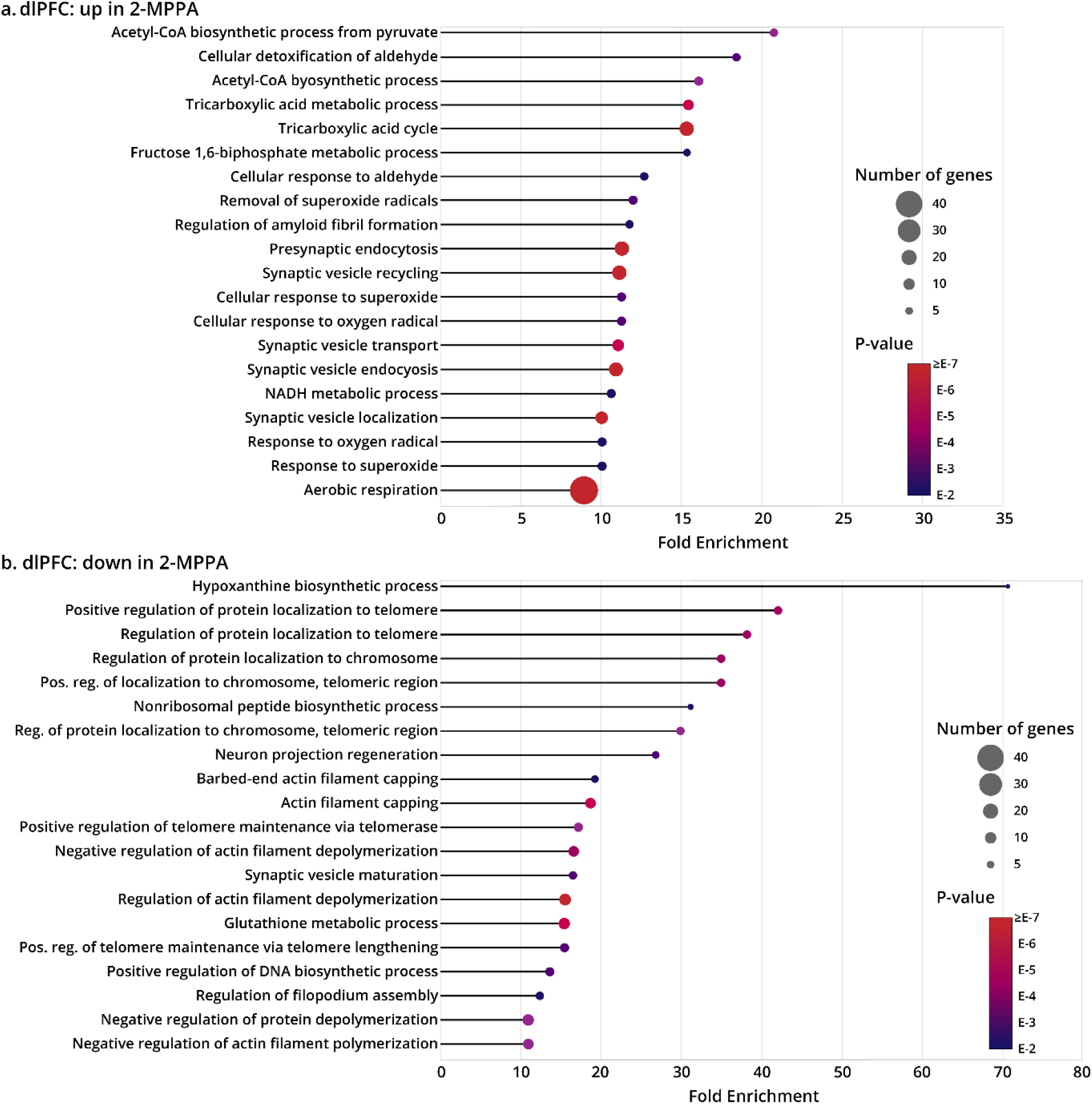
Gene ontology analysis of dlPFC proteins (**a**) upregulated or (**b**) downregulated with 2-MPPA treatment. Input protein lists, generated from label-free quantification analysis, were sorted according to both significance (p-value) and magnitude of difference (fold-change). Rank order proteins lists were analyzed using GO enrichment analysis for biological processes in *Macaca mulatta*, with the Bonferroni correction applied. The top 20 hits in either direction are represented.

**Supp. Fig. 3.**
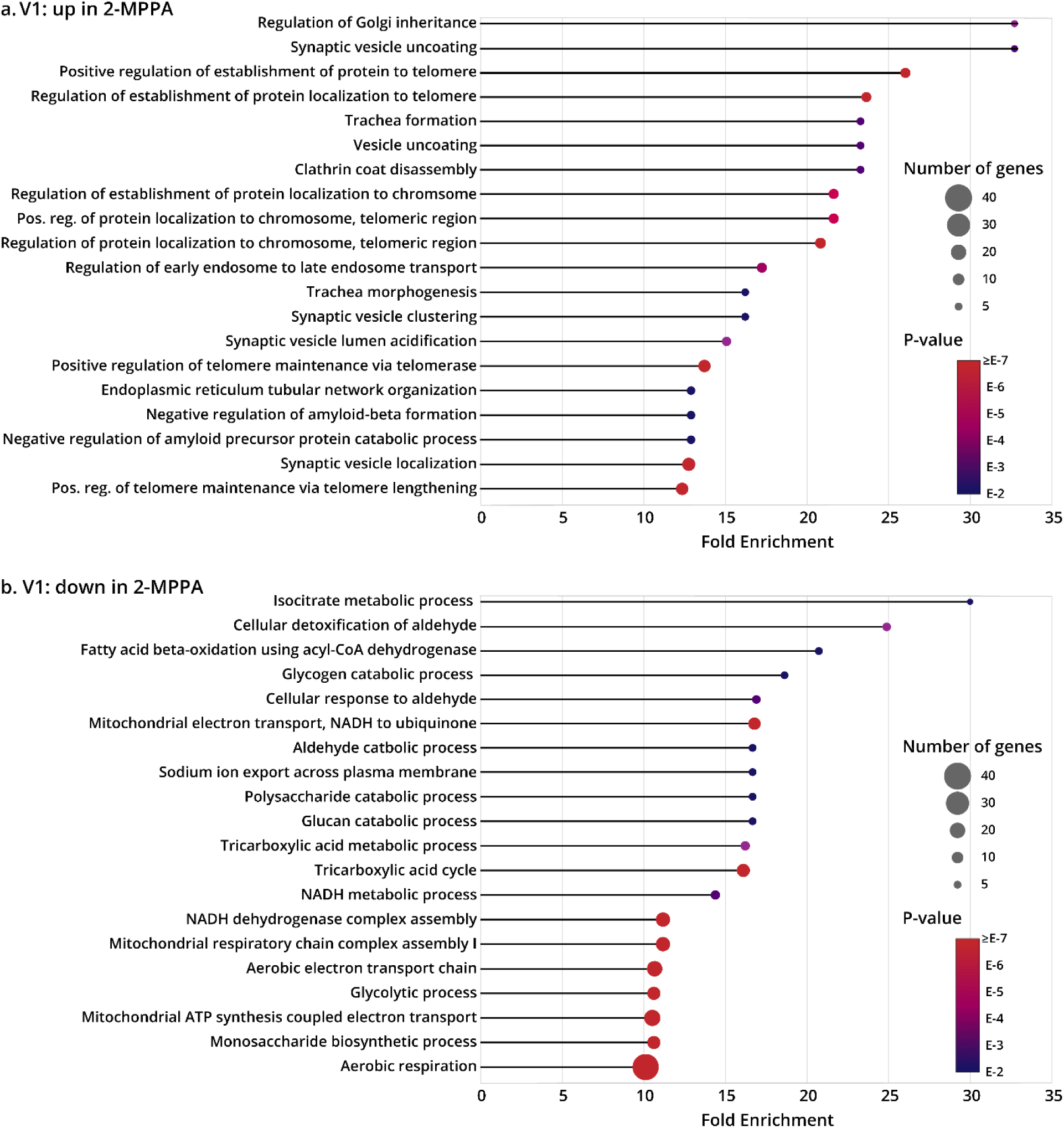
Gene ontology analysis of V1 proteins (**a**) upregulated or (**b**) downregulated with 2-MPPA treatment. Input protein lists, generated from label-free quantification analysis, were sorted according to both significance (p-value) and magnitude of difference (fold-change). Rank order proteins lists were analyzed using GO enrichment analysis for biological processes in *Macaca mulatta*, with the Bonferroni correction applied. The top 20 hits in either direction are represented.

## Acknowledgements

This study was possible due to the invaluable efforts of Michelle Wilson, Lisa Ciavarella, Sam Johnson, Tracy Sadlon, Caroline Zeiss, Jennifer Asher, Daniel Holden, Gordon Terwilliger, Alvaro Duque, and Jon Arellano. The authors would like to acknowledge the following funding sources: R21 AG079145-01, KL2 TR001862, Alzheimer’s Association Research Grant AARGD-23-1150568, and P30AG066508 Developmental Project Award (DD), and RO1 grant AG061190-01 (AFTA). ADS is supported by the National Institutes of Health Chemical Biology Training Grant (T32 GM067543).

## Conflict Of Interest Statement

Johns Hopkins University and BSS have patents US9737552B2, US20180338910A1, WO2018094334A1, and WO2016022827A1 related to GCPII inhibitors.

## Author Contributions

Conceptualization and experiments were carried out by A.D.S., A.S.B., S.A.M., and D.D. Data analysis was performed by A.D.S. & I.R.L. B.S.S. and C.H.vD. provided resources. Writing was done by A.D.S. & D.D. Writing-review and editing was performed by A.D.S., D.D., and S.A.M. The study was supervised by A.F.T.A., S.A.M., and D.D. Funding and supervision was carried out by D.D. and S.A.M. Approval of the final version of the paper was carried out by all authors.

